# Purine nucleotides are competitive inhibitors of apo-GOT1

**DOI:** 10.1101/2025.09.17.676921

**Authors:** Narges Pourmandi, Arjun Jha, Kendall Muzzarelli, Hyunsu Lee, Zahra Assar, Gordon J. Murray, Joseph H. LaPointe, Thomas P. Roddy, Gianna Medeiros, Shomit Sengupta, Costas A. Lyssiotis

## Abstract

The malate-aspartate shuttle (MAS) plays a key role in cellular metabolism by transferring electrons from cytosolic NADH into the mitochondrial matrix, thereby supporting oxidative phosphorylation, in addition to the citric acid cycle and amino acid metabolism. Here, we sought to identify allosteric regulatory metabolites of the MAS enzymes cytosolic glutamic-oxaloacetic transaminase 1 (GOT1) and mitochondrial GOT2. Using the Atavistik Metabolite Proprietary Screening platform, we identified several structurally similar metabolite hits— most notably deoxyadenosine monophosphate (dAMP) and deoxyguanosine monophosphate (dGMP)—as candidate interactors with GOT1. Follow-up thermal shift assays revealed that dAMP and dGMP destabilize GOT1 in the absence of its cofactor, pyridoxal 5’-phosphate (PLP), but have no destabilizing effect when PLP is present. Crystallographic analysis confirmed that dAMP and dGMP bind in the PLP pocket of GOT1, suggesting competitive binding. Together, these results indicate that nucleotide metabolites can interact with GOT1, offering potential insights into MAS regulation and therapeutic intervention strategies.

## Introduction

The malate-aspartate shuttle (MAS) is an essential metabolic pathway responsible for transferring electrons from cytosolic NADH into the mitochondrial matrix. NADH is membrane impermeable. Thus, to move the reducing potential in NADH into the mitochondrial matrix, this reducing power is stored in malate, which is subsequently imported into the mitochondria. In this way, the MAS plays a critical role in linking glycolytic NADH to mitochondrial oxidative phosphorylation (**Fig. 1A**). The MAS is composed of two pyridoxal 5’-phosphate (PLP) dependent aspartate transaminases (cytosolic GOT1 and mitochondrial GOT2) and two malate dehydrogenases (cytosolic MDH1 and mitochondrial MDH2). Beyond the primary function of these enzymes in redox homeostasis, they also make important products that support the citric acid cycle and amino metabolism. Rewiring of this shuttle has been observed in various cancers, where it supports high anabolic demands, highlighting its potential as a therapeutic target.^1–4^

**Figure 1.**
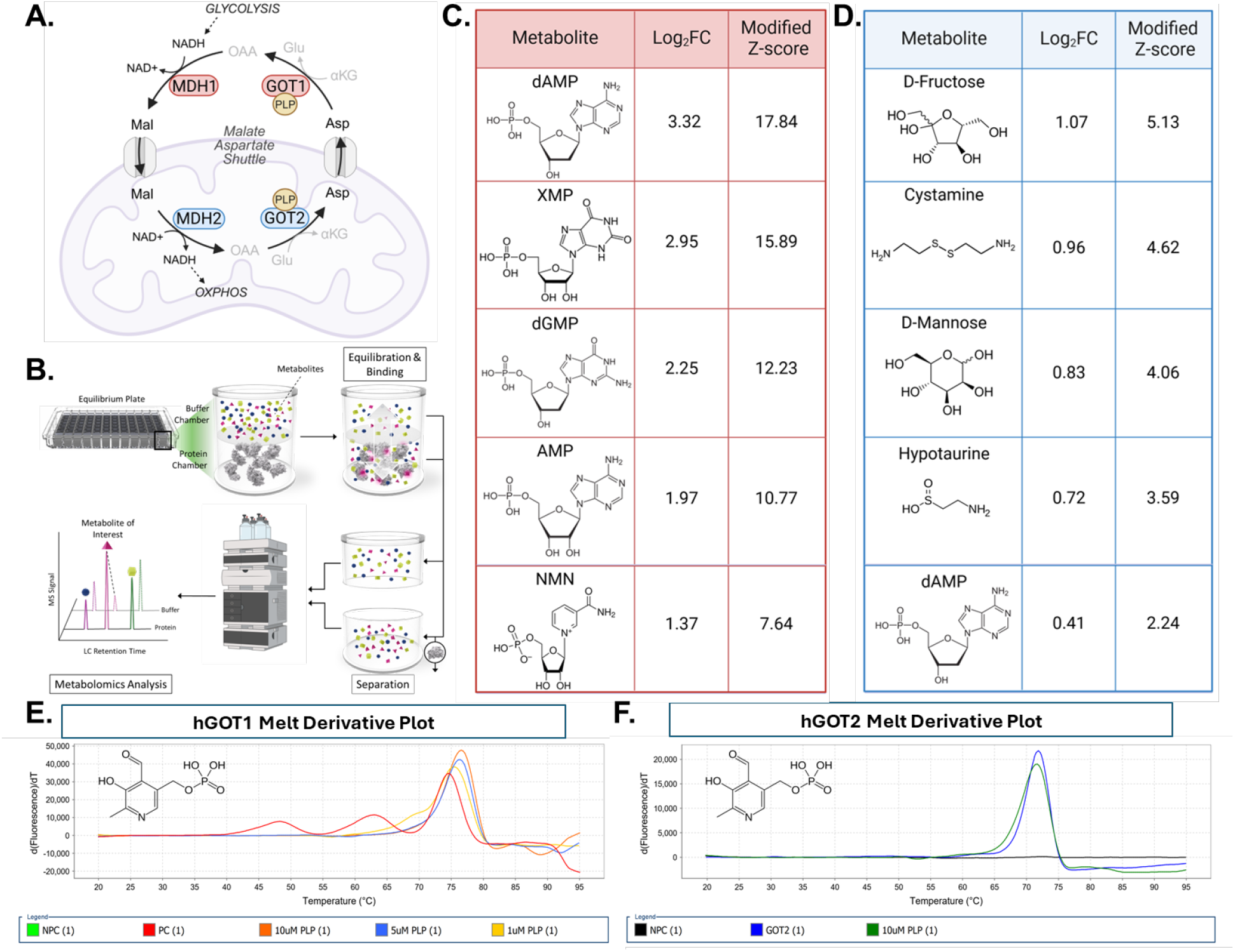
AMPS screen identifies purine nucleotides as top hits for purified GOT1 with unsaturated PLP binding sites. (**A**) The MAS moves reducing equivalents from NADH made during glycolysis onto malate for transport into the mitochondria for oxidative phosphorylation (OXPHOS). Mal: malate; Asp: aspartate; Glu: glutamate; OAA: oxaloacetate; αKG: α-ketoglutarate; PLP: pyridoxal 5′-phosphate. (**B**) Schematic of the Atavistik Metabolite Proprietary Screening (AMPS) platform. (**C**) Five of the top hits from the AMPS screen for purified GOT1. The log_2_ fold change (log_2_FC) represents the amount of metabolite found on the side where GOT1 is present over the amount of metabolite found on the side of the membrane without GOT1, where a positive log_2_FC represents more metabolite where GOT1 is present and a negative log_2_FC represents less metabolite. dAMP: deoxyadenosine monophosphate; XMP: xanthosine monophosphate; dGMP: deoxyguanosine monophosphate; AMP: adenosine monophosphate; NMN: mononucleotide. (**D**) Five of the top hits from the AMPS screen for purified GOT2. (**E**) The derivative of the melting curve of GOT1 alone (red curve) and with 1µM (yellow curve), 5µM (blue curve), or 10µM (orange curve) supplemental PLP (structure shown in left upper corner), with peaks representing the melting temperature (T_m_). NPC: non-protein control. (**F**) The derivative of the melting curve of GOT2 alone (blue curve) and with 10µM (green curve) supplemental PLP (structure shown in upper left corner), with peaks representing the T_m_.

Despite the functional importance of the MAS, little is known about allosteric, protein-metabolite regulatory interactions with the enzymes that compose the shuttle. This is largely due to the challenges of studying the dynamic and transient interactions between proteins and the metabolome. For example, the low abundance and transient nature of many metabolites make identifying interactions technically difficult. To address this technical shortcoming, Atavistik Bio developed the Atavistik Metabolite Proprietary Screening (AMPS) platform, adapted from the Mass Spectrometry Integrated with Equilibrium Dialysis for the Discovery of Allostery Systematically (MIDAS) technology.^5^ Briefly, the AMPS platform works by separating the protein of interest from a large set of metabolites by a semipermeable membrane that only allows the metabolites to diffuse between the two compartments. After equilibrium is reached, the metabolites that interact with the protein of interest accumulate on the protein side, where they can then be detected by mass spectrometry.

Leveraging this platform, we set out to characterize the metabolite-binding profiles of two MAS enzymes: cytosolic glutamic-oxaloacetic transaminase 1 (GOT1) and its mitochondrial isoform, GOT2. Through these efforts, we identified purine nucleotides, deoxyadenosine monophosphate (dAMP) and deoxyguanosine monophosphate (dGMP) as prominent interactors. Follow-up thermal shift assays, functional assays and crystallographic analysis confirmed that dAMP and dGMP bind in the PLP pocket of GOT1, suggestive of competitive binding. This work provides new insight into the metabolic relationships of GOT1 and GOT2.

## Results

The human cDNA for *GOT1* and *GOT2* were expressed in *E. Coli* and purified (**Supplementary Fig. 1**).^6,7^ We then used these proteins in the AMPS platform, screening a library containing over 500 metabolites (**Fig. 1B**). After setting a threshold of a log_2_ fold change of ≥1 to identify metabolites that had greater saturation on the protein-side, three of the top metabolites for GOT1 were deoxyadenosine monophosphate (dAMP), deoxyguanosine monophosphate (dGMP), and nicotinamide mononucleotide (NMN) (**Fig. 1C, Supplementary Table 1**). Notably, these hits share marked structural similarity with the essential GOT1 cofactor, pyridoxal 5’-phosphate (PLP). Importantly, these metabolites differed from the top metabolites for GOT2 (**Fig. 1D, Supplementary Table 2**), despite both enzymes utilizing PLP as a cofactor.

To further evaluate metabolite-protein interactions, thermal shift assays (TSA) were utilized to detect binding of metabolites to the protein through changes in protein stability. In the absence of supplemental metabolites, the melting profile of GOT1 displayed multiple peaks in the derivative fluorescence plot, with major peaks at 47.5, 62.5, and 75 °C (**Fig. 1E**). In contrast, the melting profile of GOT2 showed one major peak at 72 °C (**Fig. 1F**). The multiple peaks observed for GOT1 were hypothesized to reflect multiple oligomeric states. We later subsequently demonstrated that this was a consequence of incomplete PLP binding. Namely, when the TSA for GOT1 was run with supplemental PLP, there was one major peak at 77 °C (**Fig. 1E**). Furthermore, the melting temperature (Tm) increased proportionally with the amount of added PLP. In contrast, the melting curve for GOT2 remained unchanged upon PLP supplementation. Collectively, these findings suggest that bacterially purified GOT1, but not GOT2, harbors unsaturated PLP binding pockets.

We next assessed the impact of the identified metabolite on enzyme stability in the absence and presence of supplemental PLP. For GOT1, increasing concentrations of dAMP progressively destabilized the protein in the absence of PLP; however, this effect was not observed with PLP supplementation, with the singular melting peak remaining unaltered despite the presence of dAMP. These findings suggest that dAMP has preferential binding to empty PLP sites, destabilizing GOT1 only when these pockets are available. A similar pattern was observed with dGMP and GOT1, with dGMP leading to destabilization only when there is no supplemental PLP (**Fig. 2C-D**). For GOT2, protein stability was unaffected by dAMP under all conditions tested.

**Figure 2.**
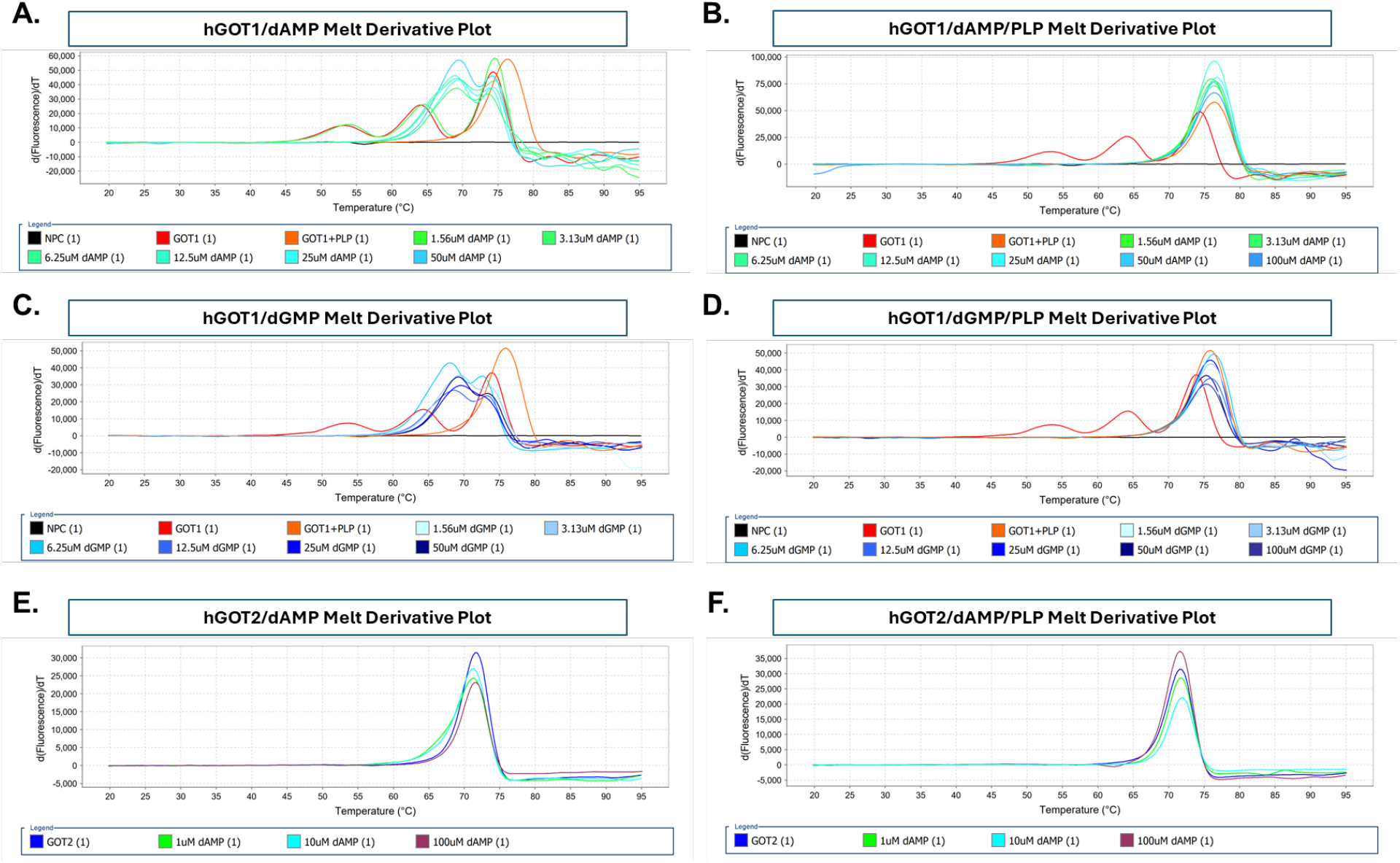
The destabilization of GOT1 by dAMP and dGMP is dependent on unsaturated PLP binding sites. (**A**) The derivative of the melting curve of GOT1 and no supplemental PLP in the presence of dAMP. For a point of comparison, GOT1 alone (red curve) and GOT1 supplemented with PLP (orange curve) are included. NPC: non-protein control. (**B**) The derivative of the melting curve of GOT1 supplemented with PLP and dAMP. (**C**) The derivative of the melting curve of GOT1 and no supplemental PLP tested with dGMP. (**D**) The derivative of the melting curve of GOT1 supplemented with PLP and dGMP. (**E**) The derivative of the melting curve of GOT2 and no supplemental PLP in the presence of dAMP. (**F**) The derivative of the melting curve of GOT2 supplemented with PLP and dAMP.

While supplemental PLP mitigated GOT1 destabilization, we hypothesized there may still be an effect on enzyme activity. We employed a reaction-coupled assay utilizing aldehyde oxidase 1 (GLOX) and horseradish peroxidase (HRP) for fluorescent monitoring (**Fig. 3A**). No significant change in GOT1 activity was observed across a range of dAMP:GOT1 molar ratios up to 1,000:1 (**Fig. 3B**). Prolonged preincubation of 24 hours of dAMP with PLP-saturated GOT1 also failed to impact activity (**Fig. 3C**). A reduction in GOT1 activity was identified only with dAMP:GOT1 ratios exceeding 1,600:1 with prolonged incubation (**Fig. 3D**), suggesting that only supraphysiological concentrations of dAMP compromise GOT1 function even when the enzyme is PLP-bound.

**Figure 3.**
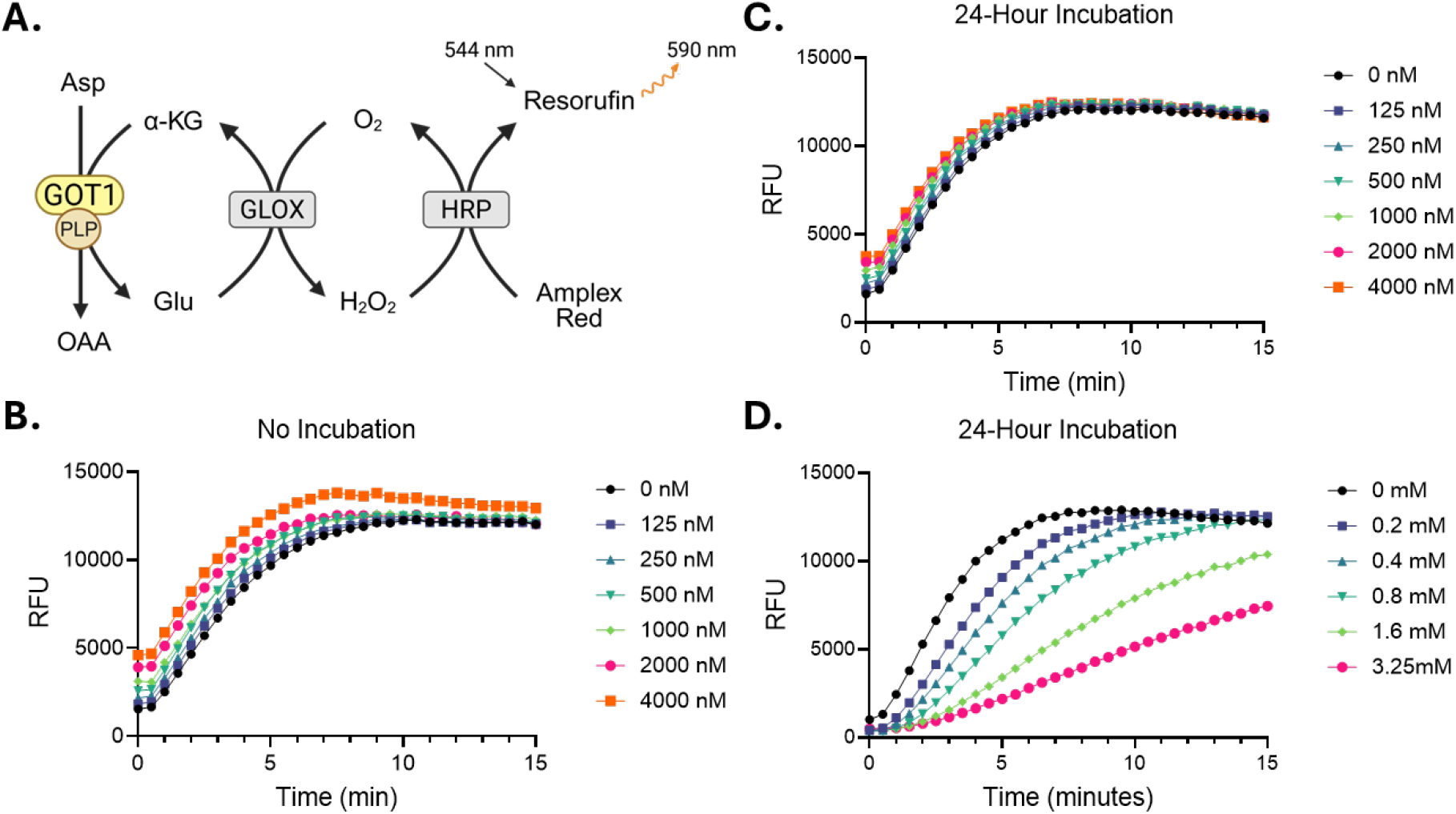
dAMP does not impact the function of PLP-supplemented GOT1 at physiological concentrations. (**A**) The reaction-coupled fluorescent reporting system utilized as the functional assay for GOT1 where the product, resorufin, can be excited at 544 nm and emission of 590 nm can be recorded. PLP: pyridoxal 5’-phosphate; Asp: aspartate; OAA: oxaloacetate; Glu: glutamate; α-KG: α-ketoglutarate. (**B**) The reaction progress of GOT1 over time immediately following the addition of PLP and dAMP. RFU: relative fluorescence units. (**C**) The reaction progress of GOT1 over time following a 24-hour incubation period with PLP and dAMP. (**D**) The reaction progress of GOT1 over time following a 24-hour incubation period with PLP and dAMP at 1,600x or greater ratios to GOT1.

Finally, to directly investigate the interaction between purine nucleotides and GOT1, crystal structures were obtained for PLP-supplemented GOT1 co-crystallized with dAMP or dGMP. In line with the previous analyses, structural analyses revealed that both nucleotides occupy the PLP-binding pocket (**Fig. 4A–D**).

**Figure 4.**
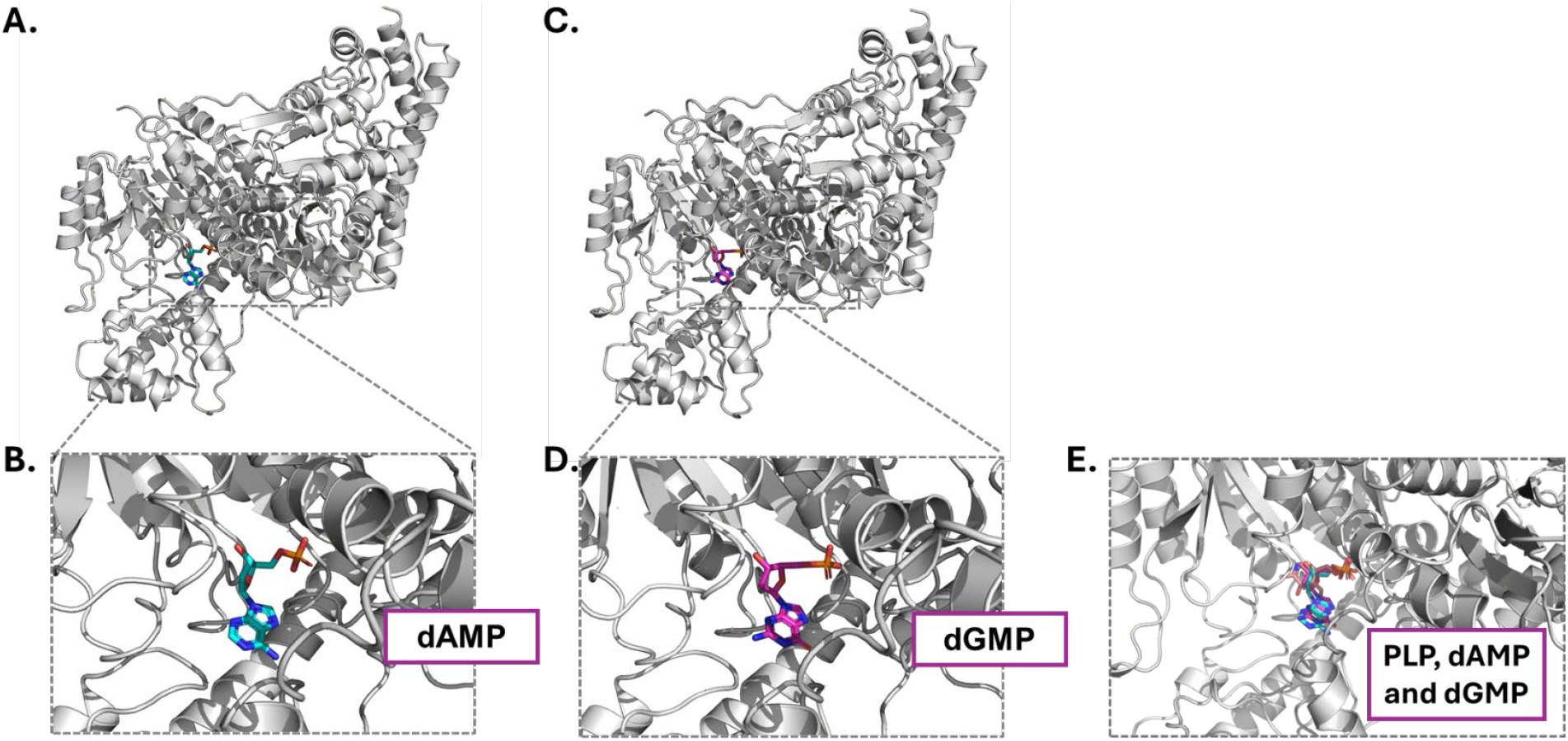
Crystal structure of GOT1 shows dAMP or dGMP in the PLP-binding site. (**A**) Ribbon representation of GOT1 (gray) in complex with dAMP (cyan). (**B**) Close-up view of the binding pocket with dAMP bound. (**C**) Ribbon representation of GOT1 (gray) in complex with dGMP (magenta). (**D**) Close-up view of the binding pocket with dGMP bound. (**E**) Overlap of PLP, dAMP, and dGMP in their binding pocket of GOT1.

## Discussion

In this study, we identify purine nucleotides as capable of binding within the PLP binding pocket of GOT1. This study was limited by the observation that supplemental PLP increases GOT1 stability, which is suggestive of unsaturated PLP sites after purification. The identification of dAMP and dGMP as possible metabolite interactors in the AMPS assay was intriguing, as purine nucleotides are not known endogenous regulators of GOT1. Given that PLP is required for GOT1 function, it was imperative to supplement follow-up experiments with dAMP and dGMP with PLP. The TSAs suggested that dAMP and PLP may compete for the same or overlapping binding sites within the enzyme—most plausibly, the canonical PLP-binding pocket, with PLP displaying higher affinity. Crystal structures confirmed the location of the binding to be within the PLP-binding pocket. Functional assays revealed that, while supraphysiological concentrations of dAMP inhibit GOT1 activity even when PLP binding pockets are fully occupied, these effects do not manifest at metabolite levels typically found in cells. Therefore, our data do not support a physiologically relevant allosteric inhibition of GOT1 by purine nucleotides under normal cellular conditions. A prudent next step would be to repeat the AMPS screen using purified GOT1 that is thoroughly saturated with PLP to more accurately capture metabolite interactions reflective of the native enzyme state. This approach could uncover subtle regulatory mechanisms masked by partial cofactor occupancy.

## Acknowledgements

We are deeply thankful to Krishnapriya Chinnaswamy and the Center for Structural Biology for their assistance in protein purification and Dr. Amy Myers for management of experimental reagents.

## Funding

This work was supported by the National Institutes of Health grants R01 CA025560 (C.A.L.), R01 CA001120 (C.A.L.), T32 GM007863 (N.P.) and T32 GM145470 (N.P.).

## Author contributions

N.P. and C.A.L. conceived the study and designed the experiments. N.P. purified the proteins, T.P.R., G.J.M. and J.H.L. performed the AMPS screen, N.P., G.M., H.L. and A.J. did the thermal shift assays, N.P. and A.J. did the functional assays, and K.M. completed the crystallization and crystal structure analysis. N.P. analyzed all the experimental data besides the crystal structures and N.P. wrote the manuscript with assistance from all authors. All authors contributed to the evaluation, integration and discussion of the results.

## Competing Interests

In the past 3 years, CAL has consulted for Astellas Pharmaceuticals, Odyssey Therapeutics, Third Rock Ventures, and T-Knife Therapeutics, and is an inventor on patents pertaining to Kras regulated metabolic pathways, redox control pathways in pancreatic cancer, and targeting the GOT1/ME1 pathway as a therapeutic approach (US patent no: 2015126580-A1, 05/07/2015; US patent no: 20190136238, 05/09/2019; International patent no: WO2013177426-A2, 04/23/2015).

## Data and materials availability

Data needed to evaluate the conclusions of the paper are present in the supplementary materials.

## Methods

### GOT1 protein purification

Human GOT1 was purified using a previously established protocol.^6^ Bacterial pellets were homogenized by being magnetically stirred at 4 °C in buffer A [20 mM NaH_2_PO_4_ (pH 7.4) and 0.5 M NaCl], 1:100 protease inhibitor, 1:100 lysozyme, and 1:10000 universal nuclease for 1 hour. Cells were lysed by pulsing sonication with a Branson model 250 sonifier at 50% power output in 3 min on/off cycles. The solution was then centrifuged at 16000 rpm for 30 min using an F20-12×50 LEX rotor at 4 °C in a Sorvall Lynx 6000 centrifuge (Thermo Scientific). The supernatant was filtered using a 0.45 μm PES filter. The clarified lysate was passed over a 5 mL HisTrap HP column (GE Healthcare) using an NGC Quest FPLC system (Bio-Rad) at a flow rate of 1 mL/min with buffer A and eluted using a 20 to 500 mM imidazole gradient over 20 column volumes (CV). The absorbance at 280 nM was monitored, and peak elution fractions were collected and concentrated to 2 mL using an Amicon Ultra-15 centrifugal filter unit with a 10 kDa molecular weight cutoff at 4000 rpm and 4 °C. Material was then injected onto a 16/60 Sephacryl S-200 HR size exclusion column (GE Healthcare) in buffer B [20 mM HEPES (pH 7.5) and 200 mM NaCl]. Dimeric material was separated from tetrameric material via SEC to ensure that proteins were monodisperse. Protein was concentrated to 10−20 mg/mL, dialyzed into 150 mM ammonium acetate pH 7.4 buffer, and flash-frozen in liquid nitrogen for storage until use.

### GOT2 protein purification

Human GOT2 was purified using a previously established protocol.^7^ The DNA plasmid was transformed into chemically competent E. coli strain BL21 (DE3) by heat shock for protein expression. A single colony of E. coli BL21 (DE3), harboring the expression vector in 30 mL of Luria-Bertani (LB) medium containing 100 μg/mL ampicillin, was cultivated at 37ºC until the optical density (OD600) reached

0.6. The cells were harvested by centrifugation at 4,000 × g for 10 min and resuspended in 3L fresh LB medium containing 100 μg/mL ampicillin. Subsequently, protein expression was induced with 1 mM isopropyl β-D-1-thiogalactopyranoside (IPTG) for 20 h at 16ºC. The cells were harvested by centrifugation at 8,000 × g for 15 min and washed with buffer A. After centrifugation, the cell pellet was resuspended in 40 mL (for 1 L culture) ice-cold extraction buffer A, and lysed by ultrasonication at ice-cold temperature. The cell lysis was centrifuged at 12,000 rpm for 15 min to separate soluble (supernatant) and precipitated (pellet) fractions. The resulting supernatant was filtered with a 0.22 μm syringe filter and then loaded onto a 5 mL HisTrapTM FF crude column (GE Healthcare, Uppsala, Sweden) pre-equilibrated with buffer A. After washing the column with buffer A containing 50 mM imidazole, the target protein was eluted with buffer C (20 mM NaH_2_PO_4_, 0.5 M NaCl, 300 mM imidazole, pH 7.4). The buffer was exchanged with 20 mM Tris buffer (pH 7.5) containing 20 mM NaCl using a 5 mL HiTrapTM desalting column (GE Healthcare). The desalted sample was loaded onto a 1 mL SP SepharoseTM FF column (GE Healthcare). After washing, the column was eluted with a linear gradient of NaCl from 20 to 500 mM in Tris buffer (pH 7.5) at a flow rate of 1 mL/min. Fractions with enzyme activity were pooled and the buffer was exchanged for 20 mM Tris (pH 7.0) containing 20 mM NaCl by using a 5 mL HiTrapTM desalting column (GE Healthcare), and then the protein was concentrated to a final concentration of 5 mg/mL with a 3 kDa cut-off concentrator (Millipore). The purity of the eluted protein was analyzed by SDS–PAGE and found to be >95%.

### AMPS metabolite screening

AMPS was performed using a system similar to that described.^5^ Briefly, 30 µM of target protein in 150 mM ammonium acetate pH 7.4 was plated in triplicate on one side of a 10 kDa MWCO dialysis membrane (SWISSCI Diaplate™) with 300 µL metabolite/compound pool aliquots in 150 mM ammonium acetate pH 7.4 placed on the reverse side. There were 8 metabolite/compound pools with ∼120 compounds per pool at an assay concentration of ∼700 nM/compound. The assay plates were sealed with adhesive aluminium foil (Thermo AB0626) and incubated at 4 °C with shaking (500 rpm) for 20 hrs. Post-dialysis protein and metabolite/compound dialysates were recovered and diluted/quenched 1:5 in 2:2:2:1:0.1 (acetonitrile:methanol:isopropylalcohol:water:formic acid), incubated at −20 °C for 1 hour, centrifuged at 3000 x g for 5 mins, with the supernatant being transferred to a 96-well PCR plate (Axygen, PCR96HSC), sealed with a silicone adhesive film (Analytical Sales and Services Inc., 96104) and placed at 4 °C in the autosampler for LC-MS analysis. The LC-MS system used is an UHPLC-QToF with an Agilent 1290 Infinity II binary pump, autosampler and column compartment (Agilent, Santa Clara, CA, USA). The UHPLC was interfaced to a Sciex 7600 QToF (Sciex, Framingham, MA, USA) operating in either electrospray positive or negative ionization mode. Pools 6 through 7 were analyzed using reversed phase chromatography (Thermo, Accucore 2.1 x 50 mm C18, 2.6 µm) using a 5 min linear gradient of 0.1% formic acid in water (mobile phase A) and 0.1% formic acid in acetonitrile (mobile phase B). Data was collected using negative electrospray in TOF MS mode for pool 6, while pool 7 was collected using positive electrospray. Pools 1 through 5 and 8 were analyzed using hydrophilic interaction liquid chromatography (HILIC) (Water, Atlantis Premier, BEH Z-HILIC, 2.1 x 50 mm, 1.7 µm) using a 5 min linear gradient of 10 mM ammonium formate in 90:10 (acetonitrile:water) pH 9.0 + 0.1% deactivator (Agilent, AG5191-4506) (mobile phase A) and 10 mM ammonium formate in 10:90 (acetonitrile:water) pH 9.0 + 0.1% deactivator (Agilent, AG5191-4506) (mobile phase B). Data was collected using positive electrospray in TOF MS mode for pools 1 & 2, pools 3 & 4 were collected using negative electrospray and pool 8 was collected using both positive and negative electrospray.

Raw data was analyzed using the Analytics application within the Sciex OS software (Sciex, Framingham, MA, USA) using targeted compound information for each pool. For each compound the peak area ratio between the protein chamber and metabolite chamber was calculated and converted to log2 fold change. The log2 converted ratios for all compounds were used to calculate the modified Z-score for each compound using the following equation:

mod_zscore = 0.6745((log2-fold change – median(log2-fold change)) / mad(log2-fold change))

Modified Z-scores greater or equal to 2 were considered presumptive hits.

### Thermal shift assays (TSA)

TSA were performed using the QuantStudio 6Pro instrument. For GOT1, the final reaction volume was 10 µL and contained GOT1 buffer (20 mM HEPES pH 7.5, 200 mM NaCl), SYPRO Orange dye (5X final concentration), GOT1 (1.0 mg/mL final concentration) with and without PLP (10 µM final concentration). For GOT2, the final reaction volume was 10 µL and contained GOT2 buffer (20 mM Tris-HCl pH 8.0, 150 mM NaCl, 1 mM DTT), SYPRO Orange dye (5X final concentration), GOT2 (0.1 mg/mL final concentration) with and without PLP (10 µM final concentration). Compounds were diluted in either PBS or DMSO and screened from 0-100 µM. A control with no protein was performed for all samples. A continuous temperature increase from 4.0 °C to 100.0 °C was scanned every 0.5 °C with a ramp speed of 0.1 °C per second. SYPRO Orange fluorescence was quantified with an excitation wavelength of 491 nm and an emission wavelength of 586 nm. Samples were run in triplicate. Melt temperatures were measured using the Protein Thermal Shift software.

### GOT1/2 functional assay

GOT1 and GOT2 enzymatic activity was measured utilizing two coupling reactions (**Fig. 3A**). First, GOT1/2 activity was coupled to two additional enzymes: aldehyde oxidase 1 (GLOX1) and horseradish peroxidase (HRP). For this reaction, aspartate and α-ketoglutarate (αKG) act as GOT1 substrates and are converted to glutamate and oxaloacetate. The glutamate product serves as a substrate for GLOX1, which is converted to αKG and hydrogen peroxide (H_2_O_2_). HRP uses H_2_O_2_ to oxidize nonfluorescent Amplex Red, resulting in the formation of the fluorescent product resorufin that can be readily measured by fluorescence. The final concentrations of the components of the assay were as follows: 100 mM HEPES buffer, 100 mM KCl, 80 nM GLOX, 0.05 unit/mL HRP, 15 mg/mL Amplex Red, 4mM aspartate, 0.5mM α-ketoglutarate and 125 nM GOT1 or GOT2. When preparing the assay, all components except GOT1 or GOT2 were combined and then added to a pre-warmed, black-walled clear-bottom 96-well plate. Meanwhile, the protein was prewarmed for 10 minutes in a 37 °C water bath. Once the protein was added, the assay was immediately put to read for 15+ minutes to determine enzyme activity by measuring absorbance in each well at 590 nm with an excitation wavelength of 544 nm. When testing 24 hour incubation, the metabolite of interest was combined with GOT1 and put in a 37 °C water bath for 24 hours. This was then added to a black-walled clear-bottom 96-well plate containing the remaining assay components.

A functional assay involving dAMP at higher concentrations was run, with single replicates for each final dAMP concentration: 0 mM, 0.2 mM, 0.4 mM, 0.8 mM, 1.6 mM, and 3.25 mM. The protocol described in the previous paragraph was utilized with a 24 hour incubation. Next, a functional assay involving dAMP at lower concentrations was run with single replicates for each final dAMP concentration: 0 nM, 125 nM, 250 nM, 500 nM, 1000 nM, 2000 nM, and 4000 nM. The protocol described before was utilized with no overnight incubation. This functional assay was also repeated with the 24 hour incubation.

### GOT1 crystallization

Crystallization was conducted using sitting drop vapor diffusion at 18°C, with diffraction quality crystals grown in 0.1 M Sodium HEPES pH 7.0-7.2 and 20% PEG 3,350. The purified GOT1 protein in 25 mM Hepes,1mM TCEP, 300mM NaCl, pH 7.0, 10% Glycerol buffer at 20 mg/mL was complexed overnight with 1 mM dAMP or dGMP then plated at a 1:1 ratio (protein:mother liquor). Crystals were cryo-protected in mother liquor containing 15% PEG 3,350 and 10% glycerol then flash frozen in liquid nitrogen and shipped to Diamond Light Source for data collection on the I04 beamline. Datasets were collected on GOT1/dAMP and GOT1/dGMP complexes that diffracted at 2-3Å, and the diffraction data was processed in the P212121 space group using Xia2 Dials. Molecular replacement for the data set was performed using a search model based on PDB ID:6DND in PHASER, PHENIX. The top solution was refined using restrained refinement in REFMAC, CCP4. Several rounds of refinement and model building were performed using COOT, CCP4 and PHENIX.

---

**Supplementary Figure 1:**
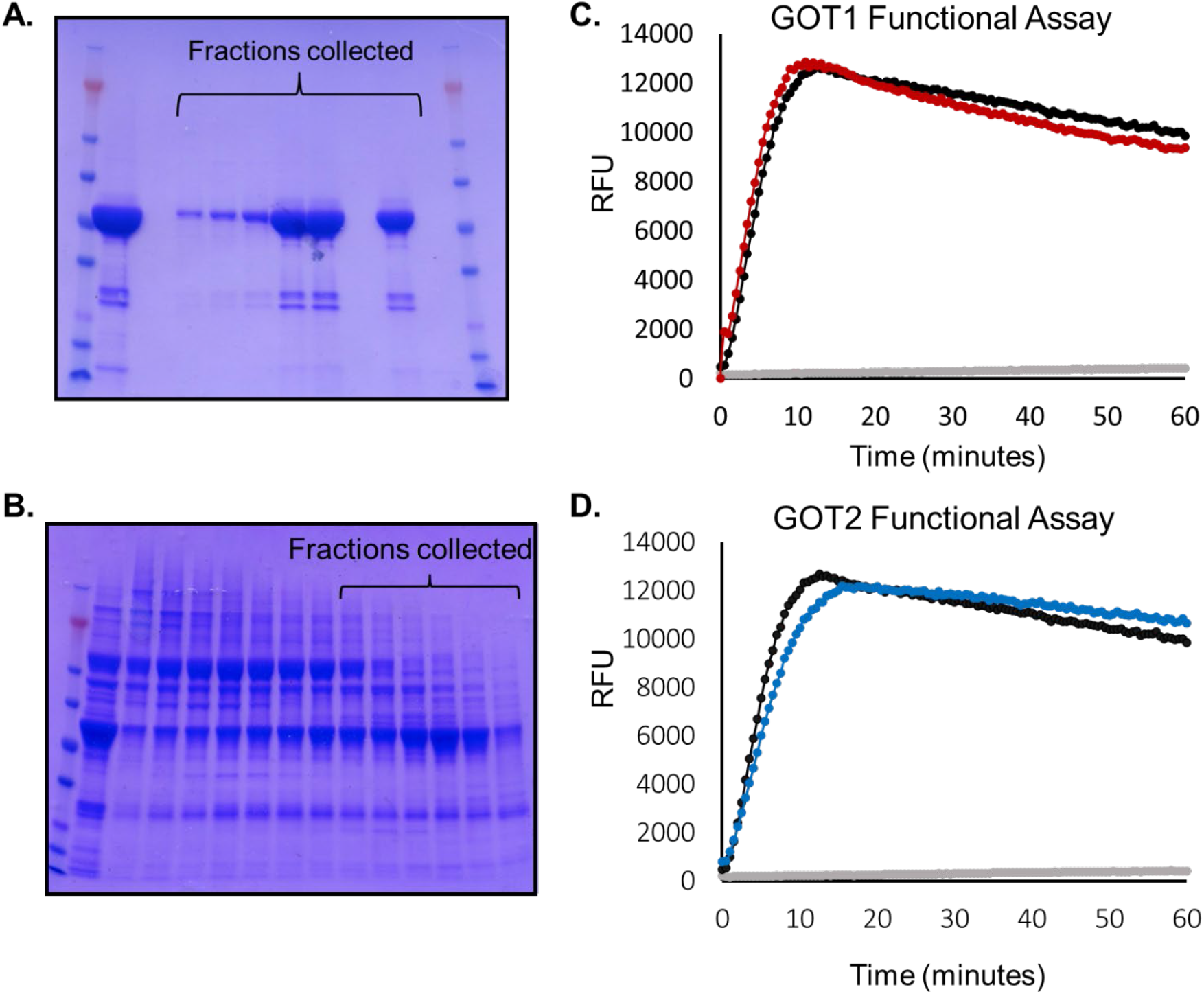
Purification and functional analysis of GOT1 and GOT2. (**A**) Protein gel displaying the fractions of the gel filtration column during the purification of GOT1 that were subsequently collected for functional assessment and screening. (**B**) Functional assay showing that the purified GOT1 (red line) is comparable in activity to a purchased GOT1 (black line). The gray line is a no protein reaction. RFU: relative fluorescence units. (**C**) Protein gel displaying the fractions of the gel filtration columns during the purification of GOT2 that were subsequently collected for functional assessment and screening. (**B**) Functional assay showing that the purified GOT2 (blue line) is comparable in activity to a purchased GOT1 (black line). The gray line is a no protein reaction.

**Supplementary Table 1:**
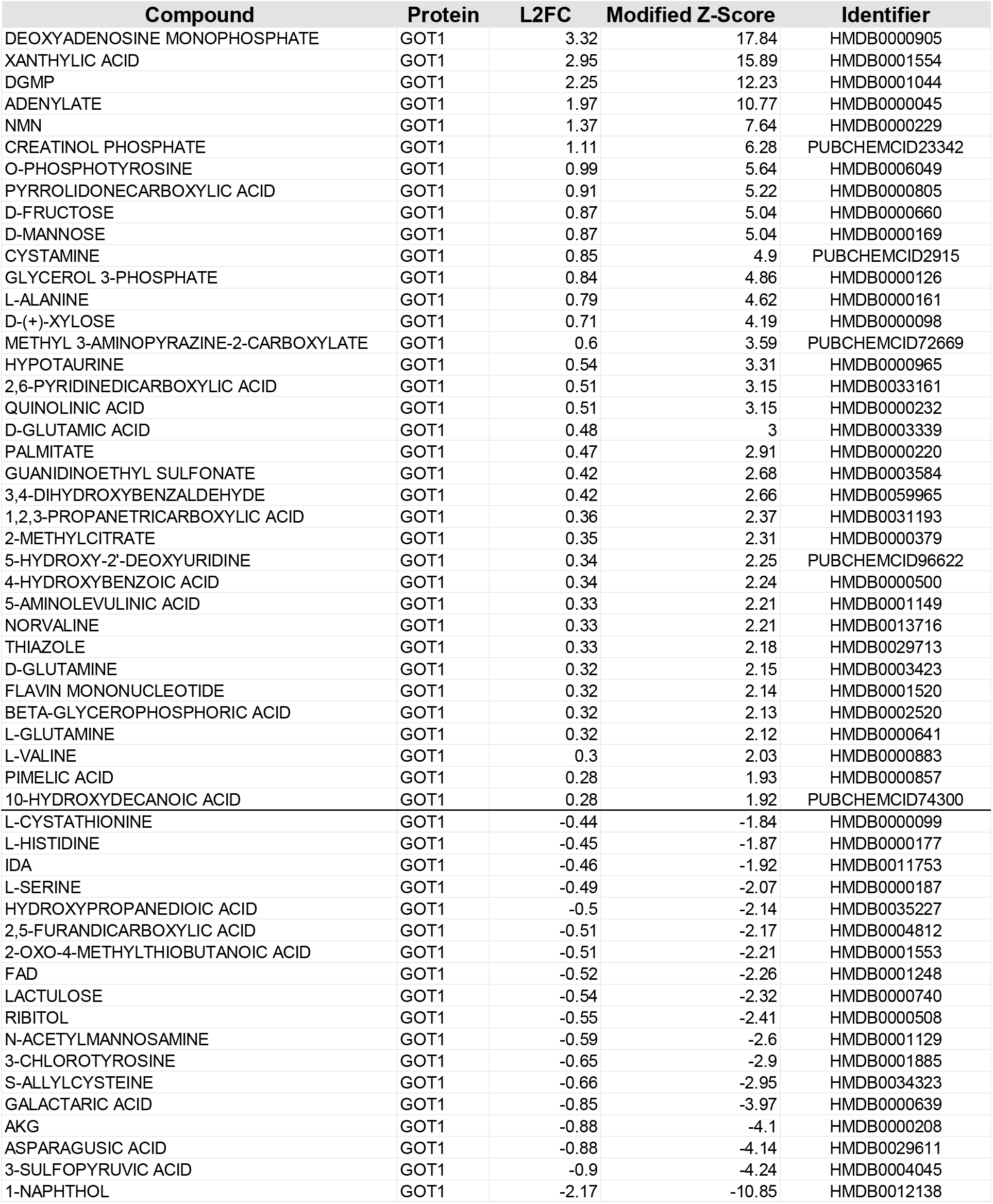
Metabolite hits from AMPS screening with GOT1.

**Supplementary Table 1:**
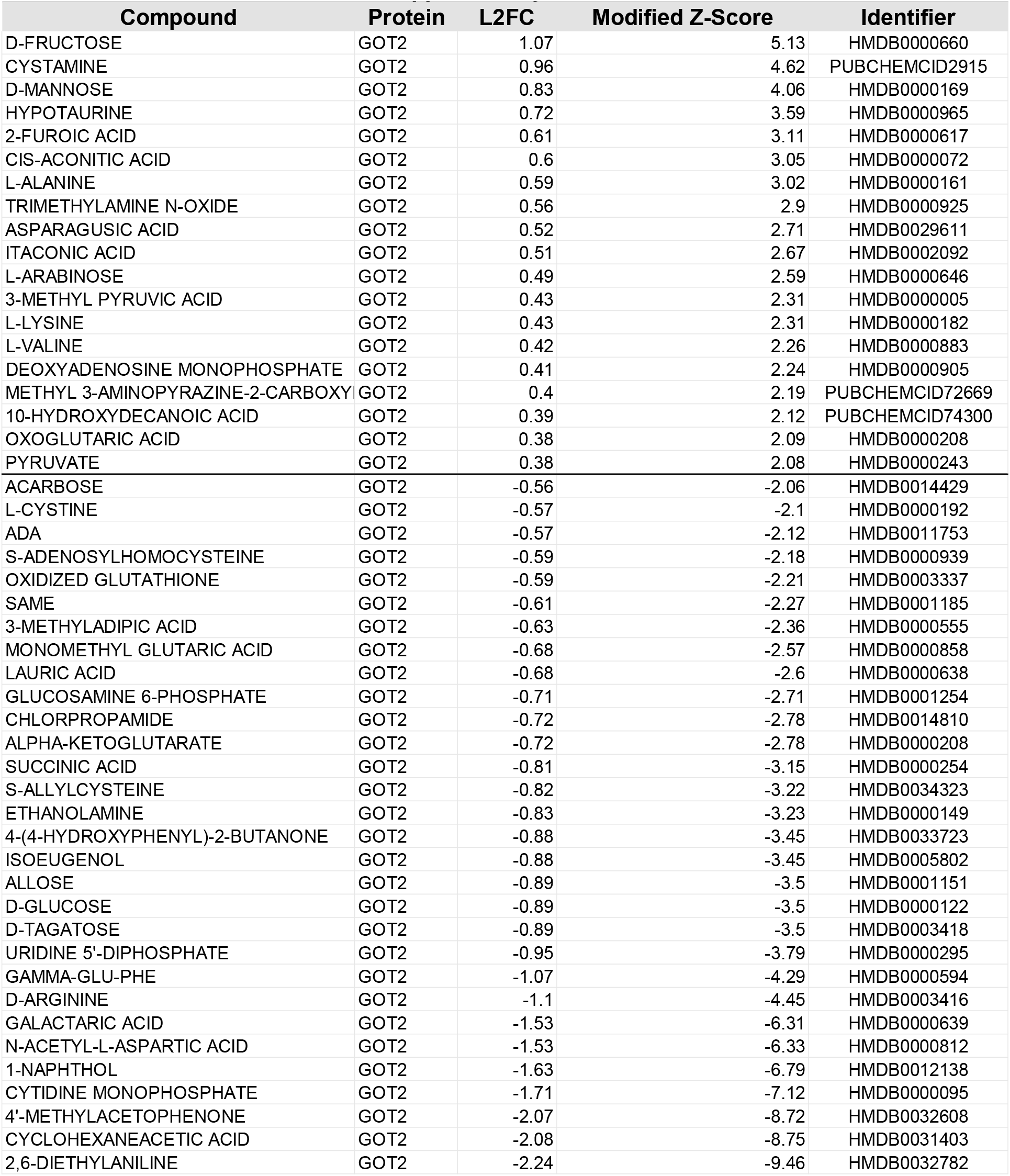
Metabolite hits from AMPS screening with GOT2.

## Notes

### Competing Interest Statement

In the past three years, C.A.L. has consulted for Odyssey Therapeutics and Third Rock Ventures, and is an inventor on patents pertaining to Kras regulated metabolic pathways, redox control pathways in pancreatic cancer, and targeting the GOT1-ME1 pathway as a therapeutic approach (US Patent No: 2015126580-A1, 05/07/2015; US Patent No: 20190136238, 05/09/2019; International Patent No: WO2013177426-A2, 04/23/2015).

